# Electrocardiographic characterization of non-selective His bundle pacing. Validation of novel diagnostic criteria

**DOI:** 10.1101/631481

**Authors:** Marek Jastrzębski, Paweł Moskal, Karol Curila, Kamil Fijorek, Piotr Kukla, Agnieszka Bednarek, Grzegorz Kiełbasa, Adam Bednarski, Adrian Baranchuk, Danuta Czarnecka

## Abstract

**Aims:** Permanent His bundle (HB) pacing is usually accompanied by simultaneous capture of the adjacent right ventricular (RV) myocardium - this is described as a non-selective (ns)-HB pacing. Our aim was to identify ECG criteria for loss of HB capture during ns-HB pacing.

**Methods:** Consecutive patients with permanent HB pacing were recruited. Surface 12-lead ECGs during ns-HB pacing and loss of HB capture (RV-only capture) were obtained. ECG criteria for loss/presence of HB capture were identified. In the validation phase these criteria and the “HB ECG algorithm” were tested by two blinded observers using a separate, sizable set of ECGs.

**Results:** A total of 353 ECG (226 ns-HB and 128 RV-only) were obtained from 226 patients with permanent HB pacing devices. QRS notch/slur in left ventricular leads and R-wave peak time in lead V6 were identified as the best features for differentiation. The 2-step HB ECG algorithm based on these features correctly classified 87.1% of cases with sensitivity and specificity of 93.2% and 83.9%, respectively. Moreover, the proposed criteria for definitive diagnosis of ns-HB capture (no QRS slur/notch in leads I, V1, V4-V6 <u>and</u> the R-wave peak time in V6 ≤ 100 ms) presented 100% specificity.

**Conclusion:** A novel ECG algorithm for the diagnosis of loss of HB capture and novel criteria for definitive confirmation of HB capture were formulated and validated. Practical application of these criteria during implant and follow-up of patients with HB pacing devices is feasible.

**Condensed Abstract:** The 2-step ECG algorithm for loss of His bundle capture based on surface ECG analysis is proposed and validated. This method correctly classified 87.1% of cases with a sensitivity and specificity of 93.2% and 83.9%, respectively.

**What’s New:**

- This is the first study that analyzes QRS characteristics during non-selective His bundle pacing in a sizable cohort of patients.
- Precise criteria and a novel algorithm for electrocardiographic diagnosis of loss of HB capture during presumed non-selective HB pacing were validated.
- QRS notch/slur in left ventricular leads was identified as a simple and reproducible feature indicating loss of HB capture or lack/loss of correction of intraventricular conduction disturbances.
- Assessment of R-wave peak time in lead V6 rather than QRS duration for diagnosis of ns-HB pacing was validated.

## Introduction

In contrast to the legacy ventricular pacing methods, His bundle (HB) pacing results in physiological activation of the ventricles via the specialized conduction system of the heart. In recent years several groups have reported encouraging outcomes of permanent HB pacing generating rapidly growing interest in this form of bradyarrhythmia and heart failure therapy.^1–9^ Permanent HB pacing is usually accompanied by simultaneous engagement of the right ventricular (RV) working myocardium near the HB - this is described as a non-selective HB (ns-HB) pacing.^10^ This new form of ventricular pacing deserves careful electrocardiograpic characterization, especially since it is present in the majority of patients who currently receive HB pacing devices. During HB pacing, high capture thresholds and significant threshold rise are observed in approximately 10% of patients.^10^ Therefore, loss of HB capture during follow-up might be relatively common and masked by the still present RV-only myocardial pacing.

Although some ECG criteria for diagnosis of ns-HB pacing were arbitrary proposed,^11^ their diagnostic value was never validated and it is currently not known if there are any ECG features/criteria that can allow conclusive diagnosis of loss of HB capture in patients with ns-HB pacing.

The aim of this study was to characterize the morphology of the QRS complex during ns-HB pacing in order to identify diagnostic features for either ns-HB capture or RV-only capture in patients implanted with HB pacing devices.

## Methods

### Study design

Consecutive patients in a tertiary cardiology center, implanted with a permanent His bundle pacing device between 2014-2019, were recruited. In all these patients permanent HB pacing was performed using a Medtronic (USA, Minneapolis) model 3830 lumenless, 4.1 French, active helix pacing lead that was screwed in to the His bundle area using standard methods for permanent HB pacing.^10, 12^ Surface 12-lead ECGs during ns-HB pacing and during loss of ns-HB capture (i.e. with RV-only capture) were recorded. We included only patients in whom QRS morphologies during ns-HB capture and RV-only capture were confirmed with differential pacing output or programmed HB pacing; these two methods served as a gold standard diagnosis.

During the exploratory phase of the study, screening of QRS features (**Figure 1**) potentially diagnostic for His bundle capture/loss of His bundle capture was conducted using a randomly selected small population (15%) of all obtained ECGs. The following ECG features were chosen for the initial analysis: 1. global QRS duration, 2. R-wave peak time in lead V6 (RWPT) and 3. mid-QRS notch or slur/plateau in leads I, II/III/aVF, aVL, V1 and V4-V6.

**Figure 1.**
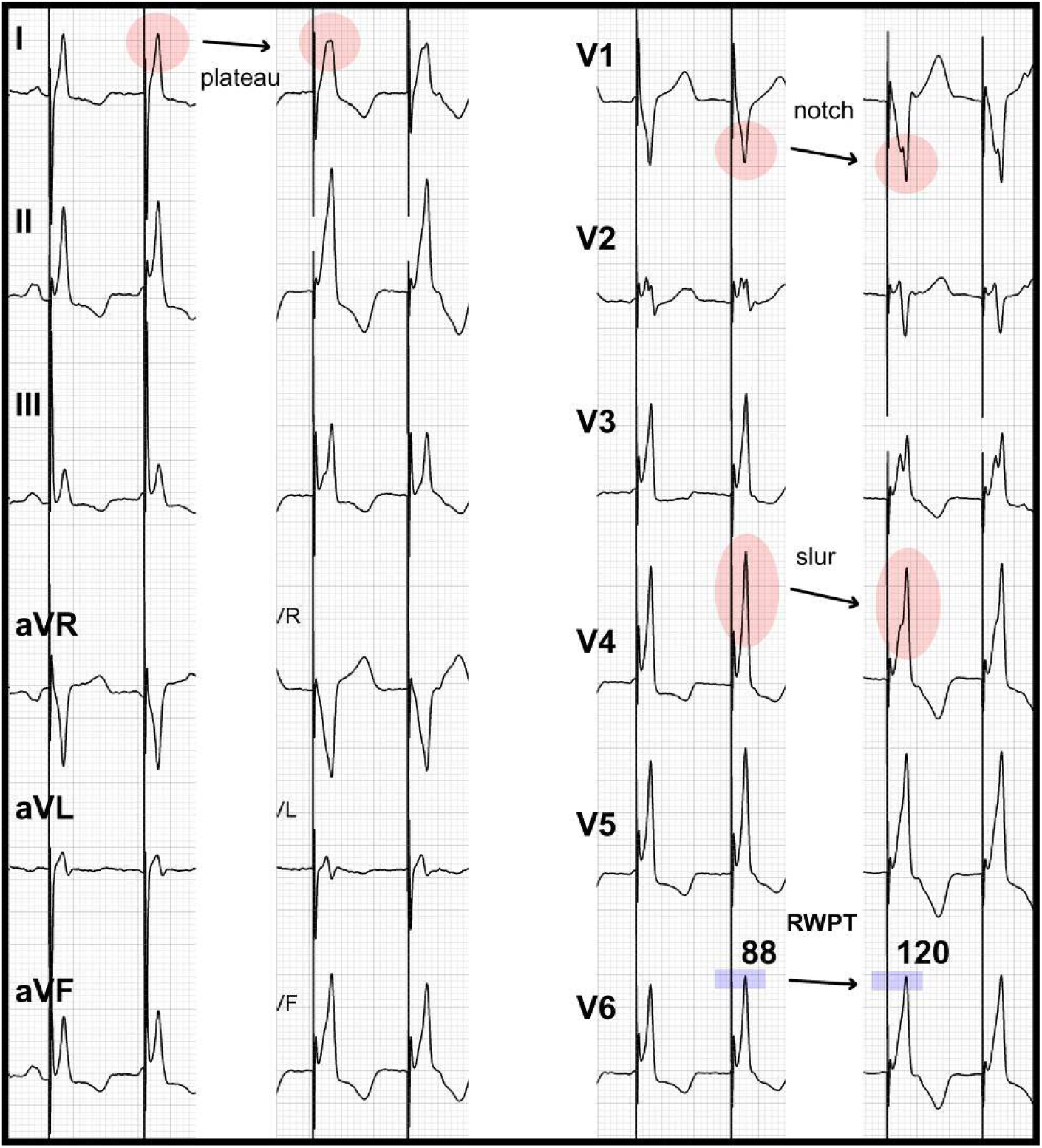
Typical changes in QRS morphology related to loss of His bundle capture in a patient with non-selective His bundle pacing. In lead I the pointy peak changes into a slur/plateau, in leads V1 and V3 a notch appears, and in leads V4 and V5 a slur develops. Global QRS duration changes from 128 ms to 162 ms and R wave peak time in lead V6 prolongs from 88 ms to 120 ms. Please note that there is a notch in the transitional QRS (V2) already during ns-HB pacing – this is why transitional QRS complexes were not included into the algorithm.

Features identified as most promising for the diagnosis of ns-HB pacing and/or RV-only myocardial capture in the exploratory phase of the study were chosen for construction of an algorithm. Moreover, criteria for definitive confirmation of HB capture were proposed. For the validation phase, we used a separate set of electrocardiograms from different patients.

During the final phase of the study, a post-hoc analysis of false positives was made. Diagnostic mistakes where ns-HB pacing was diagnosed as RV-only pacing were analyzed with the aim to elucidate the reasons behind the misleading paced QRS morphologies.

### ECG assessment

The paced QRS duration and lead V6 RWPT time were assessed in all studied ECGs using semiautomatic measurements (manually positioned digital calipers, paper speed of 100 ms/s, high signal augmentation). Duration was done according to the global QRS method (i.e., from the pacing spike to the latest QRS end in any of the 12 simultaneously recorded ECG leads), as recommended by the American Heart Association for patients with interventricular conduction disturbances and as validated in the cardiac resynchronization therapy patients CRT population.^4, 13–16^ Leads V4-V6 were assessed only if dominant R-wave was present (R or Rs with R/S ratio > 3) to avoid assessment of transitional QRS complexes. Notch was defined as two consecutive changes in the direction (≥ 90 degrees) of the R or S waves. Slur was defined as a visually evident non-gradual change of the angle of the ascending or descending slope or the R-wave (or S-wave in case of lead V1); slur angle should be between 10 - 90 degrees. Notch/slur was assessed mostly as recommended by Strauss for left bund branch block (LBBB) criteria.^17^ Additionally, we considered the top-QRS plateau/blunt peak (isoelectric or nearly isoelectric part lasting ≥ 30 ms) as a form of a slur. A criterion, that a slur must be in the upper 60% of QRS amplitude, was also introduced. This was necessary to avoid counting pseudo delta wave that is almost always present during ns-HB pacing, as mid-QRS slur.

For the validation phase of the study ECGs were printed on a millimeter paper with standard speed (25 mm/s) and augmentation (1 cm=1 mV), and saved as graphic files. Each ECG was assessed independently by two physicians (general cardiologist without any experience on HB pacing or HB ECG assessment [P.K.] and an implanting electrophysiologist with considerable experienced on HB pacing [K.C.]. Both were blinded to the established diagnosis of HB capture or loss of HB capture. In case of disagreement on ECG categorization between these two observers, consensus was reached by including a third observer.

### Statistical methods

Continuous variables are presented as means and standard deviations. Distribution of the QRS and lead V6 RWPT was estimated by the kernel method. Categorical variables are presented as percentages. Between group differences were assessed using the Fisher exact test for 2 x 2 table or Student’s t-test, as appropriate. The performance of binary decision rules was described using diagnostic accuracy (ACC), sensitivity (SN) and specificity (SP). The performance of the QRS duration and V6 RWPT in discriminating between ns-HB and RV pacing was assessed using the receiver operating characteristic curve (ROC). Statistical analyses were performed in “R”. P-values < 0.05 were considered statistically significant.

## Results

### Exploratory and validation stages

A total of 353 ECGs obtained from 226 patients were analyzed (127 patients provided both ns-HB ECG and RV-only ECG). Clinical characteristics of this cohort are presented in **Table 1**. Diagnostic properties of the ECG features that were assessed in the exploratory phase of the study (51 ECGs from 27 patients) are presented in **Table 2**. Briefly, QRS notch/slur/plateau in leads I and V4-V6 as well as lead V6 RWPT > 110-120 ms were found to be highly diagnostic for loss of HB capture while lead V6 RWPT ≤ 100 ms and no QRS notches/slurs in leads I, V1, V4-V6 were found specific for ns-HB capture. On the other hand, leads II, III, aVF and aVL were found to be not useful for the diagnosis of loss of HB capture as notch/slur was observed in these leads during both ns-HB and RV-only pacing. On the basis of these findings, a simple “HB ECG algorithm” for loss of HB capture and also criteria for a 100% definitive diagnosis of ns-HB pacing (SP of 100%) were proposed. Loss of HB capture is to be diagnosed when either there is a notch/slur/plateau in any of the leads: I, V4-V6 **<u>or</u>** V6 RWPT > 110 ms. Remaining ECGs are to be considered as most likely representing ns-HB capture. For a definitive diagnosis of HB capture during presumed non-selective pacing, the following criteria were formulated: no notches/slurs in leads I, V1, V4-V6 **<u>and</u>** RWPT is ≤ 100 ms.

**Table 1.** Basic clinical characteristics of the whole studied group (n = 226).

|  |  |
| --- | --- |
| Age [years] | 74.2 ±12.4 |
| Male gender | 155 (68.6%) |
| Comorbidities: |  |
| • Heart failure | 95 (42.0%) |
| • Coronary heart disease | 88 (38.9%) |
| • Diabetes mellitus | 82 (36.3%) |
| • Hypertension | 188 (83.2%) |
| • Serious valvular disease | 19 (8.4%) |
| LV ejection fraction [%] | 49.3 ±13.9 |
| LVEDD [mm] | 52.2 ±7.7 |
| Pacing indications: |  |
| • AV block | 63 (27.9%) |
| • SSS | 33 (14.6%) |
| • AF | 91 (40.3%) |
| • Heart failure | 39 (17.3%) |
| Intraventricular conduction disturbances: |  |
| • LBBB | 14 (6.2%) |
| • RBBB | 52 (23.0%) |
| • NIVCD | 33 (14.6%) |
| Native QRS duration [ms] | 116.9 ±25.5 |
| Baseline HV interval | 47.3 ±16.4 |

RV - right ventricular; LV - left ventricular, LVEDD - left ventricular end diastolic diameter; AV – atrioventricular, SSS – sick sinus syndrome, AF – atrial fibrillation, LBBB – left bundle branch block, RBBB – right bundle branch block, NIVCD – non-specific intraventricular conduction disturbance

**Table 2.** Exploratory phase of the study: differences in QRS characteristics between ns-HB pacing and RV pacing during loss of HB capture.

|  | **ns-HB** | **RV** | **p** |
| --- | --- | --- | --- |
|  | **(n = 27)** | **(n = 24)** |  |
| Global QRS duration [ms] | 145.3 ±16.5 | 174.8 ±26.4 | 0.000 |
| V6 R wave peak time [ms] | 96.5 ±11.5 | 129.1 ±19.0 | 0.000 |
| V6 R wave peak time ≤ 100 ms | 19 | 0 | 0.000 |
| V6 R wave peak time ≥ 110 | 2 | 23 | 0.000 |
| V6 R wave peak time ≥ 120 ms | 1 | 17 | 0.000 |
| Lead I notch/slur | 0 | 13 | 0.000 |
| aVL notch/slur | 5 | 13 | 0.010 |
| II/III/aVF notch/slur | 6 | 8 | 0.531 |
| V1 notch/slur | 0 | 7 | 0.003 |
| V4 notch/slur | 1 | 12 | 0.000 |
| V5 notch/slur | 0 | 8 | 0.001 |
| V6 notch/slur | 0 | 3 | 0.097 |
| Notch/slur in: I, V4-V6 ≥ 1 | 1 | 20 | 0.000 |
| Notch/slur in: I, V1, V4-V6 = 0 | 26 | 4 | 0.000 |

The validation phase of the study based on 302 ECGs (199 ns-HB and 103 RV-only) confirmed diagnostic usefulness of the ECG features selected during the first phase of the study and of the novel “HB ECG Algorithm” (**Table 3 and Figure 2**). Briefly, criteria for loss of HB capture had SN, SP and overall accuracy of 93.2%, 83.9% and 87.1%, respectively. Interobserver agreement on ECG categorization with the use of the “HB ECG algorithm” presented a kappa=0.817. The criteria for the definitive diagnosis of ns-HB were present in 128/199 patients with ns-HB pacing and in none with RV pacing (SN of 64.3% and SP 100%).

**Table 3.** Validation phase. Diagnostic performance of the HB ECG algorithm for the diagnosis of loss of HB capture.

|  | SN | SP | ACC |
| --- | --- | --- | --- |
| HB ECG Algorithm - observer 1 (P.K.) | 86,4% | 87,9% | 87,4% |
| HB ECG Algorithm - observer 2 (K.C.) | 94,2% | 80,4% | 85,1% |
| HB ECG Algorithm - consensus | 93,2% | 83,9% | 87,1% |
| HB ECG Algorithm - in patients with normal<br>baseline QRS and HV interval duration | 96.1%, | 96.7% | 96.5% |

HB – his bundle; SN – sensitivity; SP – specificity; ACC – accuracy; HV – His-Ventricle interval.

**Figure 2.**
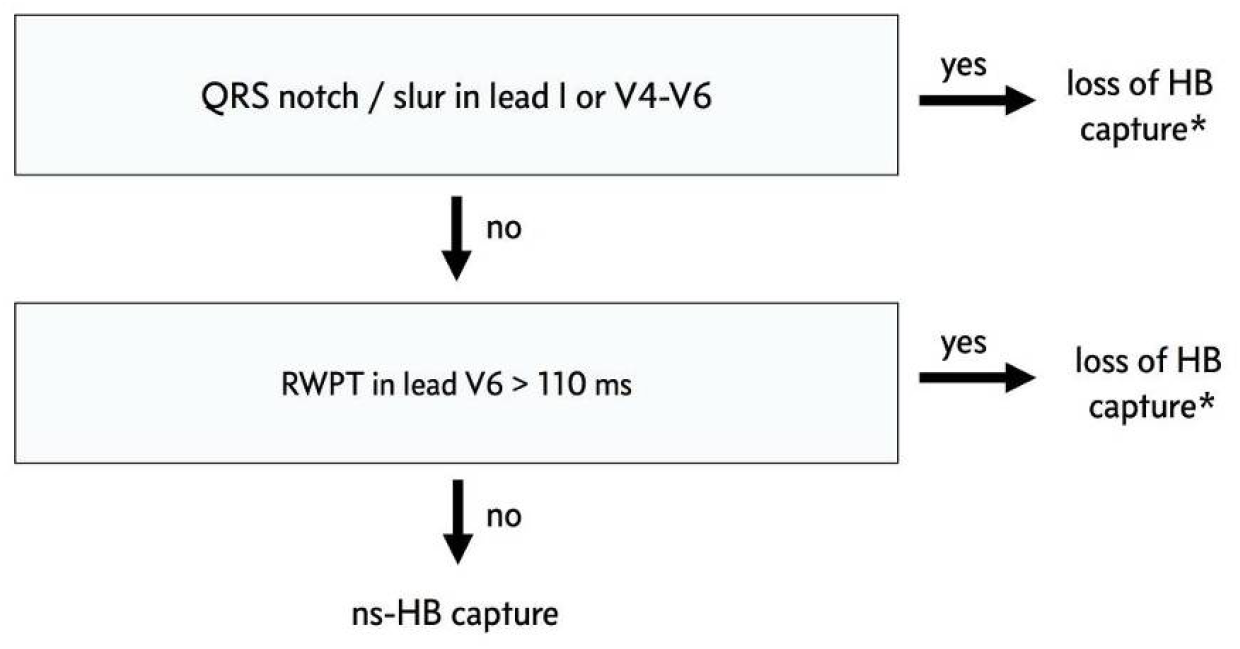
Algorithm for the electrocardiographic diagnosis of loss of non-selective His bundle capture (ns-HB). RWPT = R-wave peak time. (*) lack / loss of left intraventricular conduction disturbance correction should also be considered.

Median (quartiles) values for ns-HB and RV-only QRS duration in the whole cohort, were: 140 (132;154) ms and 172 (160;184) ms, respectively. Receiver-operating characteristic (ROC) curves for QRS duration and lead V6 RWPT calculated on the basis of the whole set of ECGs showed that V6 RWPT had bigger area under the curve (AUC) than global QRS duration (**Figure 3**). For the diagnosis of loss of HB capture, lead V6 RWPT value of 107.5 ms was identified by a ROC curve as having optimal discriminating characteristic with good balance between SN and SP of 92.1% and 86.3, respectively. Similar discriminating value for QRS duration was 151 ms with SN and SP of 90.6% and 72.1%, respectively (**Figure 3**).

**Figure 3.**
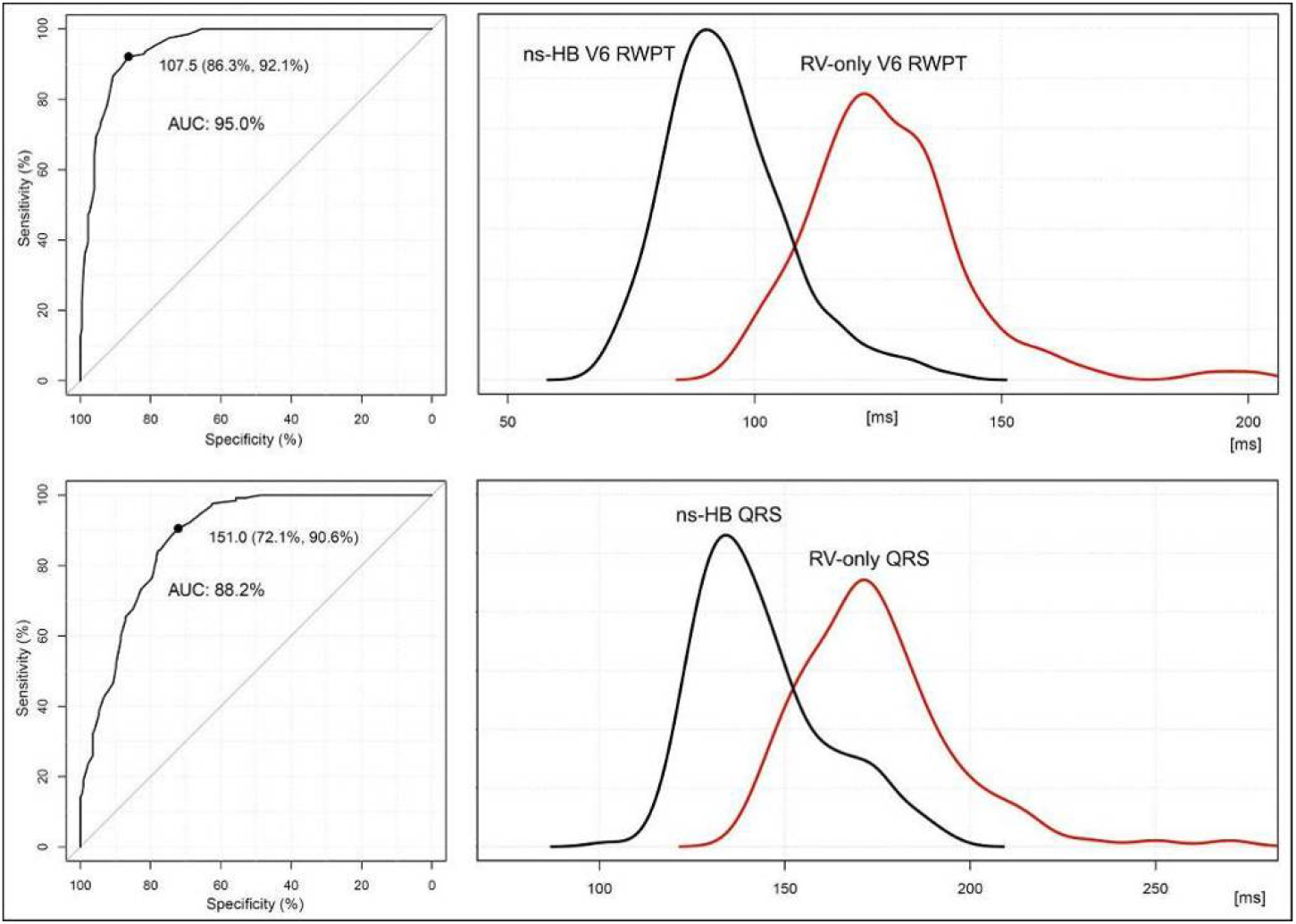
RWPT as a diagnostic criterion for loss of HB capture has greater area under the curve (AUC) than QRS duration. <u>Upper panels</u>: Receiver-operating characteristic curve for lead V6 RWPT and distribution of lead V6 RWPT values during non-selective His bundle (ns-HB) pacing and during right ventricular (RV)-only myocardial pacing. A lead V6 RWPT value close to 110 ms has best sensitivity / specificity balance for diagnosis of loss of HB capture. <u>Lower panels</u>: Receiver-operating characteristic curve for QRS duration and distribution of ns-HB and RV-only QRS duration. A QRS value close to 150 ms has best sensitivity/specificity balance for diagnosis of loss of HB capture.

### Post-hoc analysis

Post hoc analysis of patients in whom ECGs were categorized incorrectly as loss of HB capture despite confirmed HB pacing (36 cases in the whole cohort) revealed longer baseline HV interval (46.0 ms vs. 54.1 ms; p=0.006) and high percentage (86.1%) of intraventricular conduction disturbances. These false positives were caused by notch/slur in leads I or V4-V6 in 20 patients, and/or RWPT > 110 ms in 21 patients (**Figures 4 and 5**). In patients with false positive result and RWPT >110 ms, the baseline HV interval was prolonged in 42.9% of cases. The intraventricular conduction disturbances observed in these patients included: non-specific intraventricular conduction disturbances (NIVCD) in 17, LBBB in 3, right bundle branch block (RBBB) in 3, RBBB with left anterior fascicular block in 5 and RBBB with left posterior fascicular block in one, and isolated left anterior fascicular block in 2. Percentage of patients with NIVCD/LBBB in this subgroup was higher than in patients with correct diagnosis of ns-HB capture, 55.5% vs. 14.1%, respectively (p=0.000). The percentage of patients with RBBB in these two subgroups, 25.0% vs. 23.1%, respectively, did not differ.

**Figure 4.**
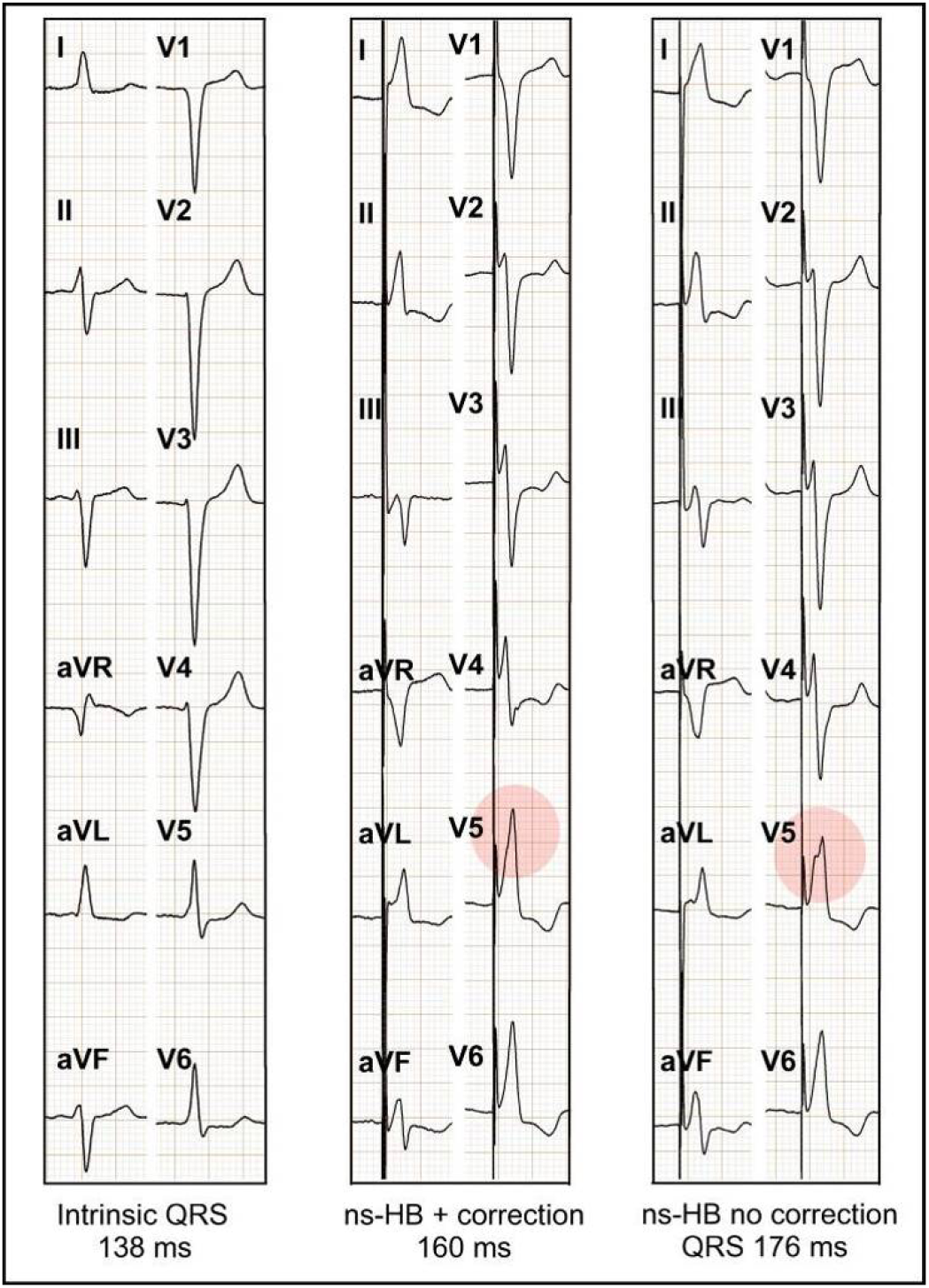
In patients with non-specific intraventricular conduction disturbances (panel A) paced ns-HB QRS often resembles QRS typically observed during right ventricular myocardial capture only (panel C). With high output pacing (panel B) there is some correction of conduction disturbances as evidenced by QRS shortening by 20 ms and disappearance of notch/slur in lead V5.

**Figure 5.**
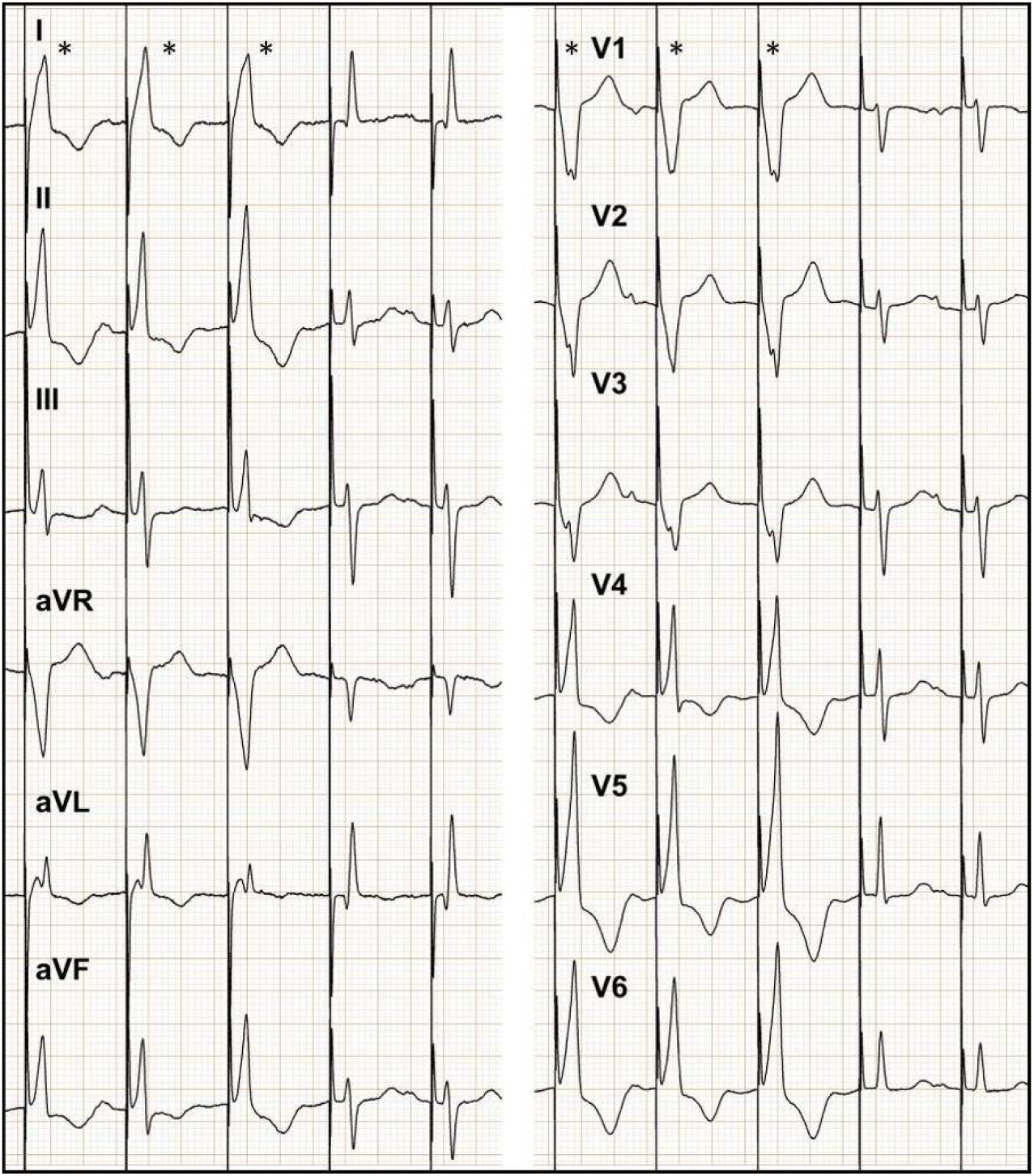
In patients with prolonged His-Ventricle (HV) interval that is not corrected with pacing, non-selective HB QRS complexes (marked with asterix) often have features typical for right ventricular myocardial capture only. In this example, HV interval of 78 ms (evidenced by stimulus to QRS interval in the last two beats, which are selectively paced HB QRS complexes), results in paced ns-HB QRS of 160 ms and lead V6 R-wave peak time of 120 ms. Importantly, even slightly bigger impact of conduction via His-Purkinje system evident in the second beat (it is a fusion complex, note the preceding P-wave) results in disappearance of the notch in lead V2 and slur in I and shortening of the R wave peak time in lead V6.

These results prompted sub-analysis of the algorithm performance in patients with normal HV interval (≤ 55ms) and normal QRS duration (<110 ms). There were 103 patients with such characteristics that have provided 143 ECG (52 RV-only and 92 ns-HB). The sensitivity, specificity and accuracy of 96.1%, 96.7% and 96.5 %, respectively.

## Discussion

The major finding of the current study is that despite significant overlap in QRS morphology between paced ns-HB QRS and paced RV-only QRS, there are also important differences. We have found that QRS notching/slurring in the left ventricular leads and R-wave peak time in lead V6 enable accurate ECG algorithm-based diagnosis for loss of HB capture in patients with permanent HB pacing devices.

### QRS notching and slurring

The development of notches/slurs in left ventricular leads - that appear immediately with the loss of HB capture - provide a criterion that is most straightforward to assess. These notches have probably similar etiology as QRS notches/slurs seen during LBBB. Loss of HB capture and activation of the left ventricle via the spread of the depolarization wavefront through the working myocardium of the interventricular septum is parallel to the situation seen with development of LBBB. Similarly, diagnostic use of QRS notch/slur for recognizing loss of HB capture parallels the use of QRS notch/slur for the diagnosis of lack of conduction in the left bundle branch (Strauss criteria for complete LBBB).^13, 15^ Interestingly, patients with preserved HB capture that were incorrectly diagnosed as loss of HB capture were characterized by baseline left intraventricular conduction disturbances, either LBBB or most commonly - NIVCD. It is quite possible that in these cases these intraventricular conduction disturbances were not corrected by HB pacing and the left ventricular activation was not much different from the activation seen during RV-myocardial pacing - leading to the development of QRS slurring/notching (**Figure 4**). Since the very purpose of HB pacing is lost in such cases, perhaps these diagnostic mistakes of our algorithm also have some clinical value. Probably the verdict of the proposed algorithm should be seen, therefore, as differentiating between correct ns-HB pacing versus either loss of HB capture or loss/lack of correction of left intraventricular conduction disturbances (LBBB/NIVCD).

### Ventricular activation times: lead V6 RWPT and global QRS duration

Fast conduction via the specialized His-Purkinje system results in more rapid depolarization of the ventricles than during RV myocardial pacing. This is the foundation for several possible duration criteria for diagnosis of ns-HB capture/loss of capture. The lead V6 RWPT criterion parallels the recognized LBBB criterion of time to intrinsicoid deflection > 60 ms in lead V6.^4^ The important difference is that in case of ns-HB pacing, the His-ventricle (HV) interval always increases RWPT by 40-50 ms. This explains why in case of ns-HB pacing, the differentiating value for RWPT must be > 110 ms rather than > 60 ms. We believe that lead V6 RWPT is better suited for diagnosis of loss of HB capture than QRS duration: firstly, it offers better separation between ns-HB and RV-only pacing than QRS duration evaluated by the distribution of QRS and lead V6 RWPT values between this two types of pacing, and by bigger area under the ROC curve (**Figures 3 and 4**); secondly, it is better associated with the primary goal of HB pacing (physiological, fast and synchronous activation of the left ventricle). This is especially evident in patients with non-corrected RBBB, where QRS is usually prolonged due to the r’ in lead V1 while RWPT in lead V6 is not influenced and remains < 100 ms (**Supplementary Figure 1**). Thirdly, RWPT is more suitable for precise measurements with the naked eye, as R-wave peak offers a very distinct point while determination of the QRS end is prone to interobserver variability, leading to the well-known imprecision of manual QRS duration assessment.

Importantly, diagnostic mistakes of our algorithm, due to RWPT > 110 ms in patients with preserved ns-HB pacing were predominantly caused by prolonged baseline HV interval that was most likely not corrected by HB pacing (**Figure 5**). In such cases, with HV interval of 60–80 ms, the depolarization wavefront from RV-myocardial capture has enough time to cross the interventricular septum and limit the contribution of the depolarization wavefront from the His-Purkinje system. Patients identified by such QRS characteristics - pointing to not complete normalization of the left ventricular activation – might benefit from additional pacing options. It was showed that simultaneous left ventricular pacing can further shorten QRS duration in patients in whom HB pacing do not fully normalize QRS complexes

The QRS duration criterion based on an arbitrarily selected cut-off point of < 120-130 ms was proposed by others for diagnosis of ns-HB pacing.^6, 11^ However, this criterion was never validated nor substantiated by a large cohort data with precise global QRS duration measurements. QRS duration during ns-HB pacing usually equals HV interval + baseline intrinsic QRS duration. Since upper normal values of HV interval QRS are 55 ms and 110 ms, respectively, then a ns-HB paced QRS, even in the absence of any intraventricular conduction disturbances, can be as wide as 165 ms. Non-corrected intraventricular conduction disturbances can further prolong the duration of a ns-HB paced QRS. An average QRS should be expected to be around 140 ms (HV of 45 ms + QRS of 95 ms), and, indeed, the median value for ns-HB QRS in our cohort was 140 ms. It is important to note that these results present a significant overlap of QRS duration values between ns-HB paced QRS and RV-only paced QRS that in some patients are quite narrow (**Figure 3**). This parallels the situation seen during differentiation between RV-only QRS and biventricular paced QRS during cardiac resynchronization therapy.^16^ Nevertheless, QRS duration can also be used for differentiation between ns-HB pacing and RV-only myocardial capture. A diagnostically optimal differentiating cut-off point, on the basis of the ROC curve analysis and very precise global QRS duration measurements, seems to be around 150 ms; while values of < 120-130 ms are 100% specific for ns-HB capture but lack sensitivity for ns-HB capture diagnosis.

### Criteria for firm diagnosis of ns-HB capture

The proposed algorithm categorizes paced ECGs in patients with presumed ns-HB pacing with adequate accuracy and compares well with ECG-based diagnostic methods from other clinical areas.^18^ However, during the implant, criteria for achieving the procedural endpoint (i.e. HB capture) must be nearly 100% definitive to serve the purpose. For this reason, we sought to additionally develop criteria that would be 100% specific for diagnosis of ns-HB pacing. These were based on absence of <u>any</u> QRS features typical for RV-only myocardial pacing i.e. lack of any notches/slurs in leads I, V1, V4-V5 or rigorously defined delayed time to R-wave peak in lead V6 (not < 110 ms but ≤ 100 ms). The validation phase confirmed that such criteria are possible and are not only 100% specific but are also surprisingly sensitive (64% of ns-HB cases). We believe that these criteria might be used as a ancillary tool during the implant to confirm HB capture, especially in cases when HB and RV capture thresholds are equal and/or in facilities without capabilities of an electrophysiological laboratory where programmed His bundle pacing might be difficult to perform.

## Limitations

The single-center recruitment of patients might have led to some bias that could reflect in ECG characteristics. The proposed criteria/algorithm might have different diagnostic value in populations with dissimilar clinical profile e.g. heart failure patients with LBBB.

## Conclusions

Novel criteria and an ECG algorithm for diagnosis of HB capture/loss of HB capture in patients with permanent ns-HB pacing were proposed and validated. The ECG algorithm might be useful during follow-up and the criteria for definitive confirmation of ns-HB capture might offer a simple and reliable ancillary procedural endpoint during HB device implantation.

---

**Supplementary Figure 1.**
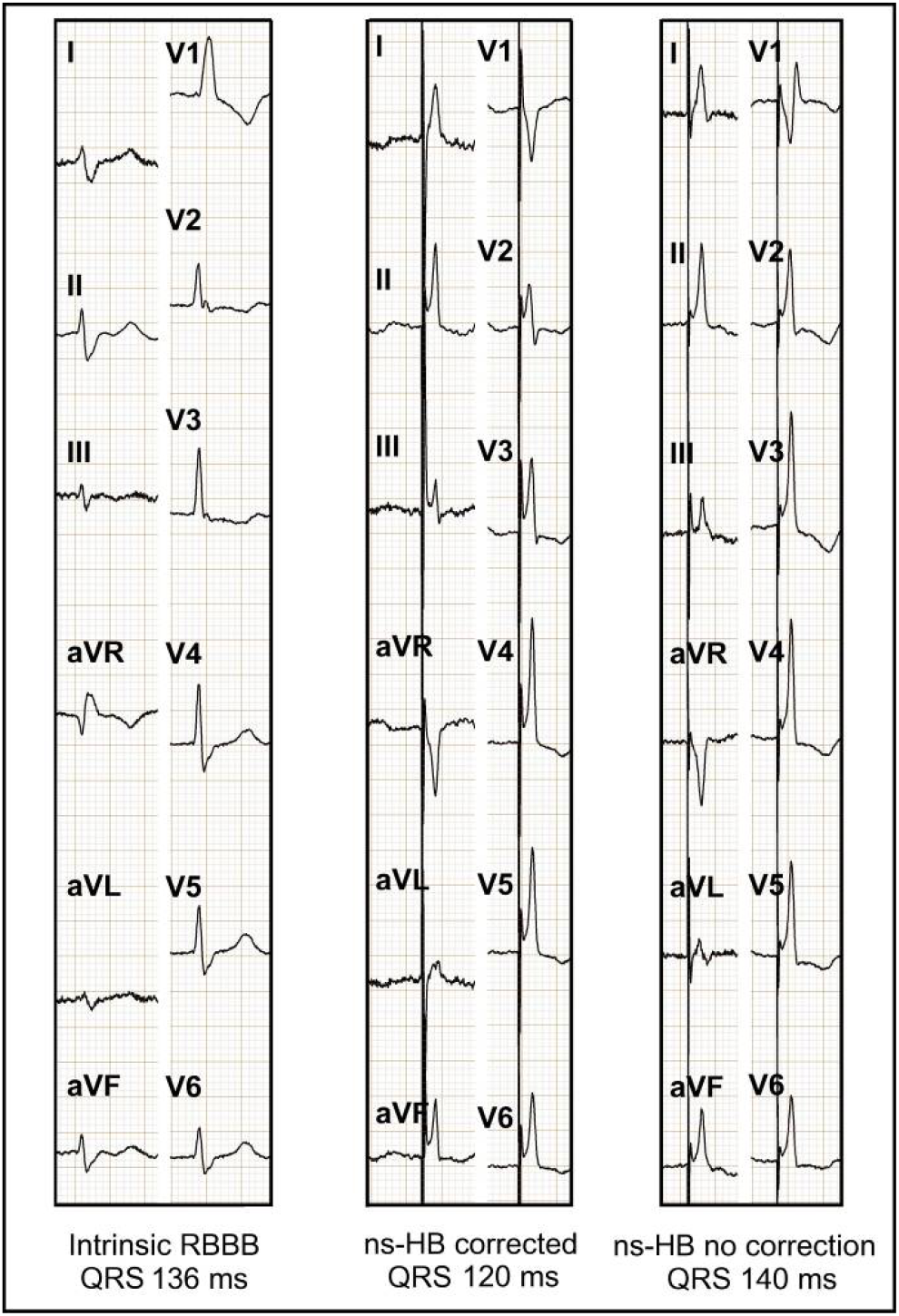
Loss of right bundle branch block (RBBB) correction during non-selective His bundle pacing (ns-HB) pacing results in QRS prolongation from 120 to 140 ms. However, the lead V6 R-wave peak time of 80 ms is not influenced and remains constant. Panel A: intrinsic QRS; panel B: ns-HB pacing with RBBB correction; panel C: ns-HB pacing without RBBB correction.

## References

1. Jastrzebski M, Moskal P, Bednarek A, Kielbasa G, Czarnecka D. His-bundle pacing as a standard approach in patients with permanent atrial fibrillation and bradycardia. Pacing Clin Electrophysiol 2018;41:1508–1512.

2. Abdelrahman M, Subzposh FA, Beer D, Durr B, Naperkowski A, Sun H et al. Clinical Outcomes of His Bundle Pacing Compared to Right Ventricular Pacing. J Am Coll Cardiol 2018;71:2319–2330.

3. Kronborg MB, Mortensen PT, Gerdes JC, Jensen HK, Nielsen JC. His and para-His pacing in AV block: feasibility and electrocardiographic findings. J Interv Card Electrophysiol 2011;31:255–262.

4. Surawicz B, Childers R, Deal BJ, Gettes LS, Bailey JJ, Gorgels A et al. AHA/ACCF/HRS recommendations for the standardization and interpretation of the electrocardiogram: part III: intraventricular conduction disturbances: a scientific statement from the American Heart Association Electrocardiography and Arrhythmias Committee, Council on Clinical Cardiology; the American College of Cardiology Foundation; and the Heart Rhythm Society: endorsed by the International Society for Computerized Electrocardiology. Circulation 2009;119:e235–e240.

5. Zanon F, Baracca E, Aggio S, Pastore G, Boaretto G, Cardano P et al. A feasible approach for direct his-bundle pacing using a new steerable catheter to facilitate precise lead placement. J Cardiovasc Electrophysiol 2006;17:29–33.

6. Occhetta E, Bortnik M, Magnani A, Francalacci G, Piccinino C, Plebani L et al. Prevention of ventricular desynchronization by permanent para-Hisian pacing after atrioventricular node ablation in chronic atrial fibrillation: a crossover, blinded, randomized study versus apical right ventricular pacing. J Am Coll Cardiol 2006;47:1938–1945.

7. Sharma PS, Dandamudi G, Herweg B, Wilson D, Singh R, Naperkowski A et al. Permanent His-bundle pacing as an alternative to biventricular pacing for cardiac resynchronization therapy: A multicenter experience. Heart Rhythm 2018;15:413–420.

8. Barba-Pichardo R, Morina-Vazquez P, Fernandez-Gomez JM, Venegas-Gamero J, Herrera-Carranza M. Permanent His-bundle pacing: seeking physiological ventricular pacing. Europace 2010;12:527–533.

9. Huang W, Su L, Wu S, Xu L, Xiao F, Zhou X et al. Long-term outcomes of His bundle pacing in patients with heart failure with left bundle branch block. Heart 2019;105:137–143.

10. Vijayaraman P, Chung MK, Dandamudi G, Upadhyay GA, Krishnan K, Crossley G et al. His Bundle Pacing. J Am Coll Cardiol 2018;72:927–947.

11. Occhetta E, Bortnik M, Marino P. Future easy and physiological cardiac pacing. World J Cardiol 2011;3:32–39.

12. Dandamudi G, Vijayaraman P. How to perform permanent His bundle pacing in routine clinical practice. Heart Rhythm 2016;13:1362–1366.

13. Jastrzebski M, Kukla P, Kisiel R, Fijorek K, Moskal P, Czarnecka D. Comparison of four LBBB definitions for predicting mortality in patients receiving cardiac resynchronization therapy. Ann Noninvasive Electrocardiol 2018;23:e12563.

14. Jastrzebski M, Baranchuk A, Fijorek K, Kisiel R, Kukla P, Sondej T et al. Cardiac resynchronization therapy-induced acute shortening of QRS duration predicts longterm mortality only in patients with left bundle branch block. Europace 2019;21:281–289

15. Strauss DG, Selvester RH, Wagner GS. Defining left bundle branch block in the era of cardiac resynchronization therapy. Am J Cardiol 2011;107:927–934.

16. Jastrzebski M, Kukla P, Fijorek K, Czarnecka D. Universal algorithm for diagnosis of biventricular capture in patients with cardiac resynchronization therapy. Pacing Clin Electrophysiol 2014;37:986–993.

17. Almer J, Zusterzeel R, Strauss DG, Tragardh E, Maynard C, Wagner GS et al. Prevalence of manual Strauss LBBB criteria in patients diagnosed with the automated Glasgow LBBB criteria. J Electrocardiol 2015;48:558–564.

18. Jastrzebski M, Kukla P, Czarnecka D, Kawecka-Jaszcz K. Comparison of five electrocardiographic methods for differentiation of wide QRS-complex tachycardias. Europace 2012;14:1165–71

